# Design of a multi-epitope camel mastitis vaccine candidate using a reverse vaccinology approach

**DOI:** 10.1101/2024.05.26.595936

**Authors:** Edwin Murungi, Nathan Langat, Ednah Masila, Ruth Onywera, Irene Ogali, Nimmo Gicheru, Hezron Wesonga, Monicah Maichomo

## Abstract

Camel mastitis caused by a diverse array of pathogenic bacteria including *Staphylococcus aureus* and *Streptococcus agalactiae* is responsible for enormous socioeconomic losses in the arid and semi-arid regions of Africa where camels are an invaluable source livelihood and nutrition. Although *S. aureus* and *S. agalactiae* are environmental contaminants, the latter is an infectious zoonotic agent. The development of a safe, efficacious vaccine for camel mastitis is a compelling priority particularly against backdrop of climate change and antimicrobial resistance. We have sequenced and assembled the genomes of local strains of *S. aureus, S. agalactiae, L. lactis, E. faecium* and *E. gallinarum* isolated from milk samples of pastoralists’ camel herds in Kenya, analysed the subtracted proteomes for antigenic proteins using VaxiJen and ascertained the allergenicity, subcellular localisation and the number of transmembrane helices of the putative candidates using AllerTOP, PSORTb and DeepTMHMM. Furthermore, B-cell and T-cell epitopes uncovered in the antigenic protein sequences using Bepi-Pred-2.0, NetCTL 1.2 and NetMHCII 2.3 were used to design a potential vaccine candidate that exhibited adequate immunogenicity, stability and flexibility without toxicity and allergenicity. Collectively, we have delineated a potential multi-epitope vaccine for camel mastitis.

## 1 Introduction

Dromedary camels (*Camelus dromedarius*) are resilient, multi-purpose animals well adapted for survival in harsh environments characterised by scarcity of water and pasture (Darwish, 2023; Seligsohn *et al.,* 2020). Camels are a vital source of income, milk and meat for pastoralists in Arid and Semi-Arid Lands (ASALs) of Africa (Guliye *et al.,* 2007; Volpato and Howard, 2014) with camel milk being an important alternative source of protein and a particularly appropriate functional food for infants and geriatrics due to its high nutrients content in these regions (Alhaj and Kanhal, 2010; Rahmeh *et al.,* 2022). Moreover, camel milk has been reported as a potential remedy for jaundice, tuberculosis, diabetes, asthma and leishmaniasis (Swelum *et al.,* 2021).

Mastitis, an intramammary infection characterised by physical, chemical and bacteriological alterations in milk, and pathological changes in the glandular tissue (El Tigani-Asil *et al.,* 2020), causes enormous agricultural losses due to diminished milk production, poor quality milk and microbial contamination of milk (Dzayee *et al.,* 2022). Besides hampering animal productivity and significantly constraining farming earnings, the disease also adversely affects human health (Aqib *et al.,* 2022; Seligsohn *et al.,* 2020). The infection may be clinical (acute or chronic) or sub-clinical. Broadly, clinical mastitis is characterised by hardening, swelling and pain of the mammary gland coupled with discolouration and clotting of milk. Specifically, in acute mastitis, the mammary secretions are watery, yellowish and blood-tinged while fibrosis and keratinization of the the udder tissue is prevalent in chronic mastitis (El Tigani-Asil *et al.,* 2020). On the other hand, in sub-clinical mastitis, inflammation without overt symptoms prevails (Aqib *et al.,* 2022). Clinical and sub-clinical camel mastitis diminish the yield, alter the properties, and impair the preservation and processing of milk (Kashongwe *et al.,* 2015).

Bacterial infections are the commonest cause of camel mastitis (Seifu and Tafesse, 2010). Although an array of bacterial species including *Streptococcus sp., Staphylococci sp., Salmonella sp., Enterococcus sp, Escherichia sp., Mycoplasma sp., Corynebacterium sp., Pseudomonas sp*., and *Campylobacteria sp*. have been isolated from camel udders (Zeryehun and Abera, 2017), the highly infective *Streptococcus agalactiae* (group B *Streptococcus*) is the main mastitis causing bacteria. Our recent studies (Ochieng *et al.,* 2021; Murungi *et al.,* 2022; Maichomo *et al.,* 2023) have identified several bacterial strains that potentially cause camel mastitis in Kenya. Given the indiscriminate use of antibiotics for the treatment of mastitis against the backdrop of global antimicrobial resistance (Naranjo-Lucena and Slowey, 2023), development of effective vaccines for camel mastitis is a pressing critical need.

Following sequencing and assembly of the genomes of *Staphylococcus aureus*, *Streptococcus agalactiae*, *Lactobacillus lactis*, *Enterococcus faecium* and *Enterococcus gallinarum* strains isolated from mastitis-infected pastoralists’ camels in Kenya, we undertook *in silico* analyses and elucidated several putative antigenic vaccine candidates from which we identified an array of epitopes that we used to design a potential multi-epitope camel mastitis vaccine.

## 2 Materials and Methods

### 2.1 Data retrieval

The proteomes of *S. aureus* (Accession: JARJBG010000027.1), *S. agalactiae* (Accession: JANCLS000000000), *L. lactis* (Accession: JAOPKV000000000), *E. faecium* (Accession: JAOTOH000000000 and *E. gallinarum (*Accession: [JAOTOI010000000](https://www.ncbi.nlm.nih.gov/nuccore/JAOTOI000000000.1/)) were retrieved from the GenBank. Non-redundant sequences in each proteome were obtained by clustering using CD-HIT (Fu *et al.,* 2012) at a sequence identity cut-off of 50%.

### 2.2 Selection of potential vaccine candidate protein sequences

Antigenicity was set as the core criteria in choosing potential vaccine candidates. The five bacterial proteomes were screened using VaxiJen v2.0 (https://www.ddg-pharmfac.net/vaxijen/VaxiJen/VaxiJen.html) (Doytchinova and Flower, 2007) at a cut-off value of 0.7. Thereafter, the subcellular localization of the selected sequences and presence of signal peptides to differentiate between secretory and non-secretory proteins was determined using PSORTb (Yu *et al.,* 2010) and SignalP 5.0 server (Almagro Armenteros *et al.,* 2019).

### 2.3 Physicochemical characterization of putative vaccine candidates

Transmembrane topology of the transmembrane proteins in the identified set of probable vaccine candidates was ascertained using DeepTMHMM (Hallgren *et al.,* 2022). Adhesion-like characteristics and allergenicity of the putative candidates were predicted using SPAAN (Sachdeva *et al.,* 2004) at a cut-off of 0.5 and AllerTOP server (Dimitrov *et al.,* 2014) respectively.

### 2.4 Linear B-cell epitope prediction

The selected antigenic sequences were screened for linear B-cell epitopes using Bepipred Linear Epitope Prediction 2.0 tool (Jespersen et al., 2017) in the Immune Epitope Database and Analysis Resource (IEDB, https://www.iedb.org/). The potential of exposed, non-allergenic epitopes to trigger antibody production was ascertained using IgPred (Gupta *et al.,* 2013).

### 2.5 Prediction of cytotoxic T-lymphocyte (CTL) and helper T-lymphocyte (HTL) epitopes

Probable CTL epitopes in the candidate protein sequences were uncovered using the NetCTL 1.2 server (Larsen *et al.,* 2007) using default settings (threshold = 0.75). Using artificial neural networks, NetCTL 1.2 integrates prediction of peptide major histocompatibility complex (MHC) class I binding, proteasomal C terminal cleavage and TAP transport efficiency allowing prediction of CTL epitopes restricted to 12 MHC class I supertype. Thereafter, helper T cells 15-mer epitopes were predicted using NetMHCII 2.3 server (Jensen *et al.,* 2018). Three human HLA-DR class II alleles were evaluated. According to standards, the lowest consensus scores of the peptides presumed to be the best binders have a lower percentile rank indicating higher affinity. An IC50 cut-off of ≤ 50 and percentile rank <1 were set as the selection criteria. The epitopes unveiled were further evaluated for their propensity to induce Th1 type immune response characterised by the production of IFN-gamma using the IFNepitope server (Dhanda *et al.,* 2013). In this case, 15-mer HTL epitopes were selected based on an IC50 value of ≤50 nM, lowest percentile rank score and highest prediction score which point to avidity of epitope binding (Nielsen *et al.,* 2008).

### 2.6 Construction of a putative multi-epitope camel mastitis vaccine

A multi-epitope potential vaccine was constructed using an array of the predicted CTL and HTL epitopes. The CTL and HTL epitopes were linked using AAY and GPGPG linkers respectively (Rana and Akhter, 2016; Kalita *et al.,* 2016; Pang *et al.,* 2022). The long alpha chain of human IL-12 () was included as an adjuvant to amplify the immunogenicity of the vaccine construct (Chatterjee *et al.,* 2021). The adjuvant was linked to the initial CTL epitope through an EAAAK linker at the N-terminal of the sequence. A histidine tag was included in the C-terminal of the vaccine construct.

### 2.7 Evaluation of antigenicity, allergenicity, immunogenicity and toxicity

The antigenicity of the candidate vaccine construct was evaluated using VaxiJen v2.0 server while the allergenicity was assessed using AlgPred 2.0 (Sharma *et al.,* 2021) and AllerCatPro (Maurer-Stroh *et al.,* 2019).

### 2.8 Physiochemical properties and solubility prediction

The physiochemical properties of the putative vaccine including theoretical isoelectric point (pI), estimated half-life, instability index, molecular weight (M_w_), aliphatic index and grand average of hydropathy (GRAVY) were determined using the ExPASy ProtParam tool (Gasteiger *et al.,* 2003).

### 2.9 Decoding the secondary and tertiary structure of the putative vaccine candidate

The three-dimensional (3-D) structure of the potential vaccine candidate was inferred via homology modeling on I-TASSER server (Yang and Zhang, 2015). I-TASSER identifies structural templates in the Protein Data Bank (PDB) by homology-independent multiple threading approaches and constructs 3-D models via multiple threading alignments and template-based fragment assembly simulations. A template modeling (TM) score generated quantifies the model quality. The quality of the 3-D model was ascertained using the ProSA web server (Wiederstein and Sippl, 2007).

### 2.10 Molecular docking of the vaccine construct onto TLR2

To determine the potential of the vaccine candidate to trigger innate immune responses, docking of the putative vaccine candidate onto TLR-2 (PDB ID:5D3I) was performed using the ClusPro server (Kozakov *et al.,* 2017). Toll-like receptors activate immune responses by recognising pathogen-associated molecular pattern molecules (PAMPs) on bacteria, viruses, fungi and parasites. The docked complex was visualised using PyMOL (Schrodinger and Delano, 2020).

### 2.11 Molecular dynamics (MD) simulation

To further delineate the interaction between the potential vaccine candidate and TLR-2, molecular dynamics simulations were undertaken using the iMODS server (Lopéz-Blanco *et al.,* 2011).

### 2.12 Simulation of the host immune response

Prediction of how the vaccine candidate would elicit an immune response was conducted using the C-ImmSim server which is underpinned by the Celada-Seiden algorithm that models immune reactions in a vaccinated mammalian system (Rapin *et al.,* 2010). In the simulations, apart form the assumption that the candidate vaccine would be administered without a lipopolysaccharide (LPS) adjuvant, all other parameters were at a default setting. The sigma-70 factor of RNApolymerase subunit was used as the negative control antigen with the measurement of immune heterogeneity undertaken and interpreted using the Simpson index D (Dolley *et al.,* 2023).

## 3.0 Results

### 3.1 Identification of potential vaccine candidates

Analyses of the subtracted proteomes of *S. aureus, S. agalactiae, L. lactis, E. faecium* and *E. gallinarum* using VaxiJen v2.0 revealed an array of highly scoring (>0.7) antigenic sequences (Supplementary Table 1) for which the top sequences were selected for further analyses (Table 1). Notably, in selecting sequences for further downstream analyses, we focussed on *S. aureus* and *S. agalactiae* given they are the principal etiological agents of mastitis in the field (Seligsohn *et al.,* 2020). The sequences selected as potential vaccine candidates (Table 1) included PDZ domain-containing protein, YpmS family protein, cell wall synthase accessory phosphoprotein MacP, pathogenicity island protein and VraH family protein.

**Table 1:** An array of proteins predicted to be antigenic using VaxiJen and evaluated for antigenicity, allergenicity, sub-cellular localisation, presence or absence of a signal peptide and the number of transmembrane helices.

| Bacterial source | GenBank Accession | Protein name | VaxiJen Score | Subcellular localization | Transmembrane helices | AllerTOP prediction | Signal peptide |
| --- | --- | --- | --- | --- | --- | --- | --- |
| *S. aureus* | MDF3296226.1 | hypothetical protein P3G69_09200 | 1.4155 | Cytoplasmic membrane | 1 | Probable non-allergen | No |
| *S. aureus* | MDF3345085.1 | cell surface protein, partial | 1.4003 | Extracellular | 1 | Probable non-allergen | No |
| *S. aureus* | MDF3296234.1 | hypothetical protein P3G69_09240 | 1.3336 | Cytoplasmic membrane | 1 | Probable non-allergen | No |
| *S. aureus* | MDF3296909.1 | DUF4887 domain-containing protein | 1.1962 | Cytoplasmic membrane | 1 | Probable non-allergen | No |
| *S. aureus* | MDF3294667.1 | elastin-binding protein EbpS | 1.136 | Cytoplasmic membrane | 1 | Probable non-allergen | No |
| *S. aureus* | MDF3294497.1 | hypothetical protein P3G69_00220 | 1.028 | Extracellular | 0 | Probable non-allergen | No |
| *S. aureus* | MDF3295037.1 | type I toxin-antitoxin system Fst family toxin | 0.9862 | Cytoplasmic membrane | 1 | Probable non-allergen | No |
| *S. aureus* | MDF3296810.1 | pathogenicity island protein | 0.9513 | Cytoplasmic membrane | 1 | Probable non-allergen | No |
| *S. agalactiae* | MCP9190290.1 | cell wall synthase accessory phosphoprotein MacP | 0.9461 | Cytoplasmic membrane | 1 | Probable non-allergen | No |
| *S. aureus* | MDF3295131.1 | hypothetical protein P3G69_03545 | 0.9448 | Extracellular | 1 | Probable non-allergen | No |
| *S. aureus* | MDF3343883.1 | hypothetical protein P3G70_08555 | 0.9351 | Extracellular | 1 | Probable non-allergen | No |
| *S. aureus* | MDF3294606.1 | hypothetical protein P3G69_00775 | 0.9042 | Cytoplasmic membrane | 0 | Probable non-allergen | No |
| *S. aureus* | MDF3296979.1 | VraH family protein | 0.8981 | Cytoplasmic membrane | 1 | Probable non-allergen | No |
| *S. aureus* | MDF3294509.1 | preprotein translocase subunit YajC | 0.8864 | Cytoplasmic membrane | 1 | Probable non-allergen | No |
| *S. aureus* | MDF3294841.1 | hypothetical protein P3G69_02025 | 0.8767 | Extracellular | 0 | Probable non-allergen | No |
| *S. aureus* | MDF3295053.1 | putative metal homeostasis protein | 0.867 | Extracellular | 0 | Probable non-allergen | No |
| *S. aureus* | MDF3296041.1 | hypothetical protein P3G69_08260 | 0.8409 | Cytoplasmic membrane | 1 | Probable non-allergen | No |
| *S. agalactiae* | MCP9190898.1 | YpmS family protein | 0.8236 | Cytoplasmic membrane | 1 | Probable non-allergen | No |
| *S. agalactiae* | MCP9190874.1 | PDZ domain-containing protein | 0.8131 | Cytoplasmic membrane | 0 | Probable non-allergen | No |

### 3.2 B and T cell epitope mapping

#### 3.2.1 B cell epitopes

The Bepipred Linear Epitope Prediction 2.0 tool within the IEDB server was used to determine B cell epitopes in the prioritised antigenic sequences (Table 2). Bepipred 2.0 predicts B-cell epitopes in a protein sequence using a Random Forest algorithm trained on detecting epitopes and non-epitopes in crystal structures of antibody-antigen complexes and outputs a prediction score for each amino acid in the input sequence whereby residues with scores above the threshold (default value is 0.5) are predicted to be part of an epitope.

**Table 2:**
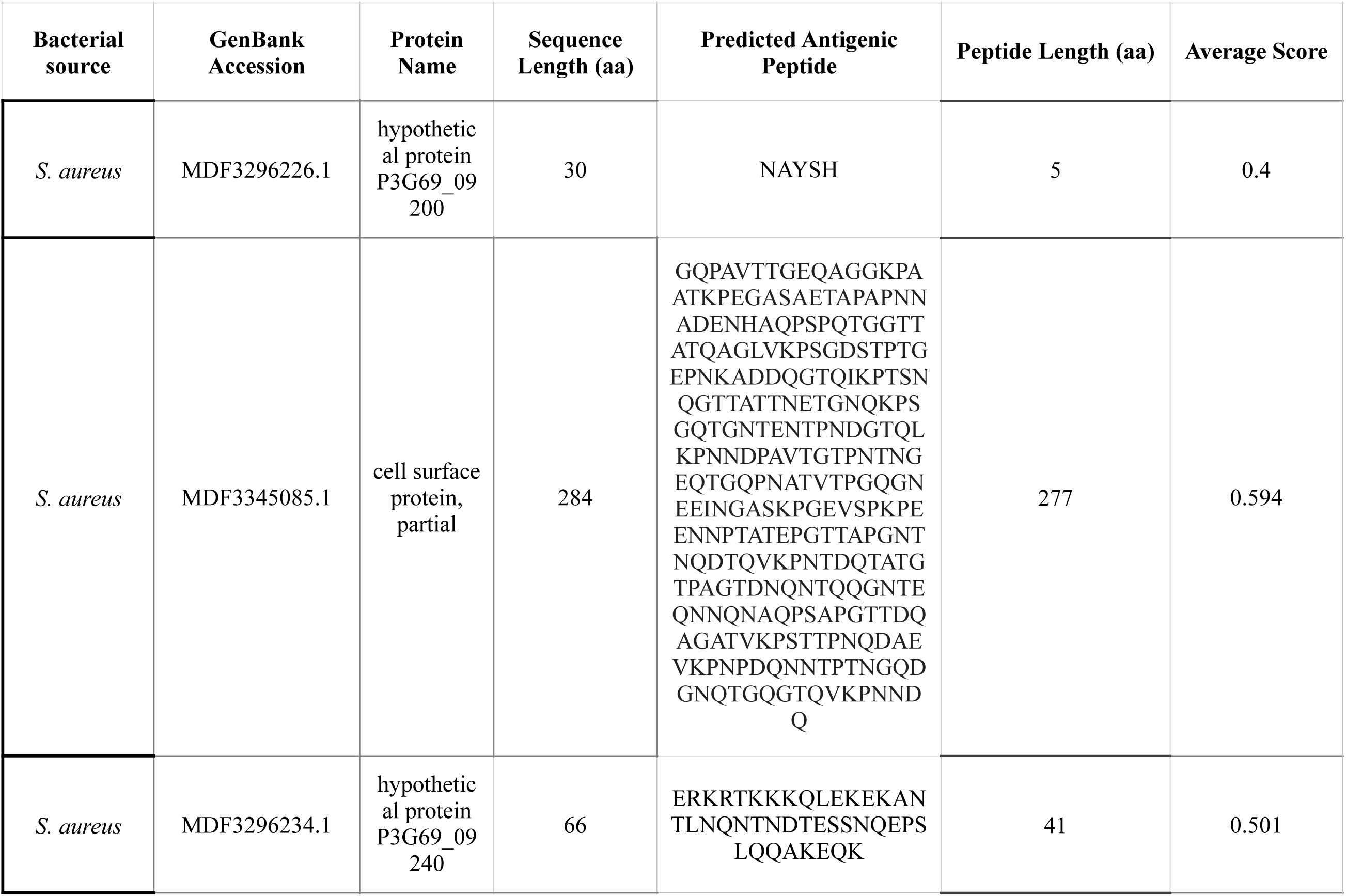
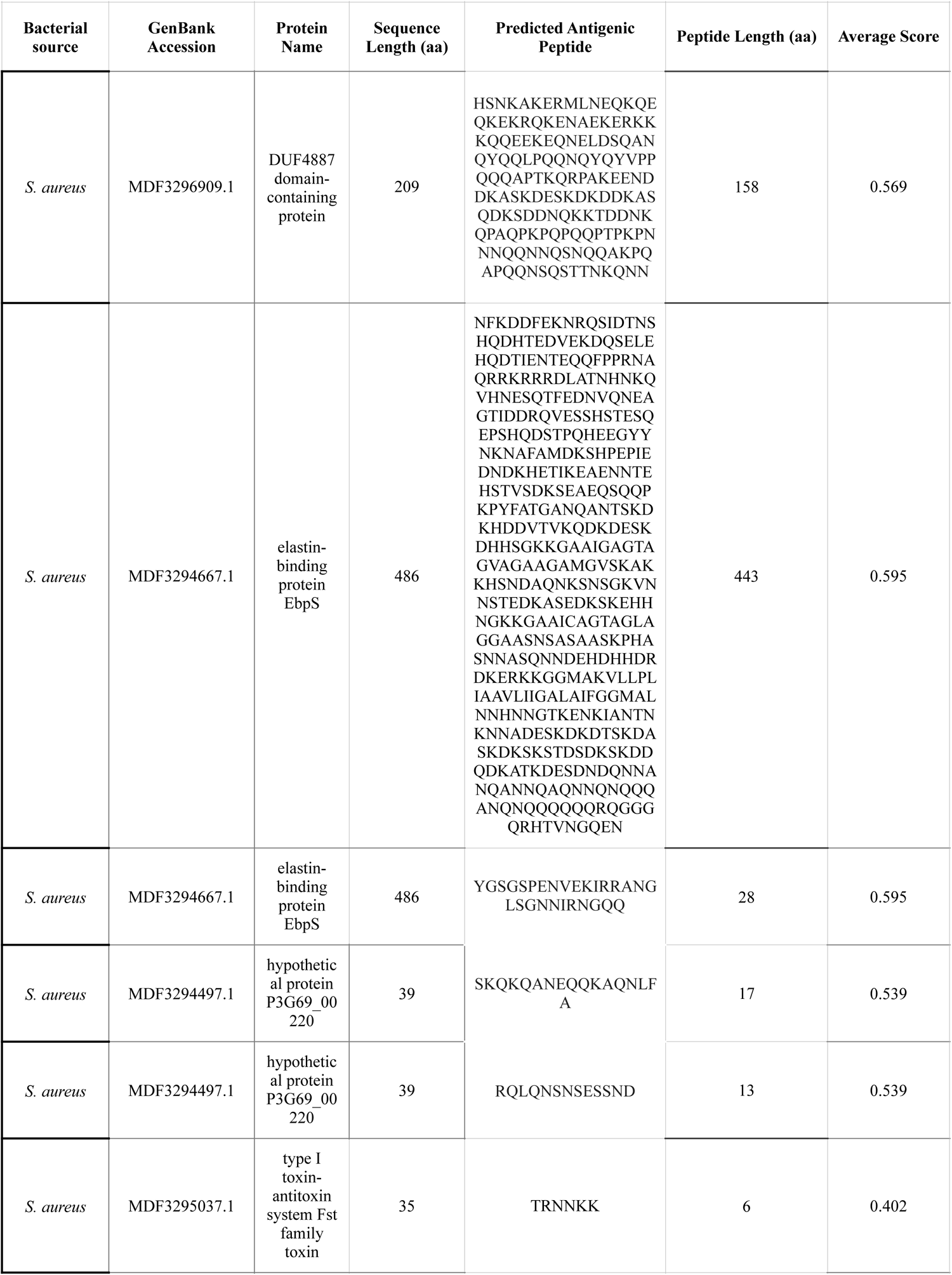
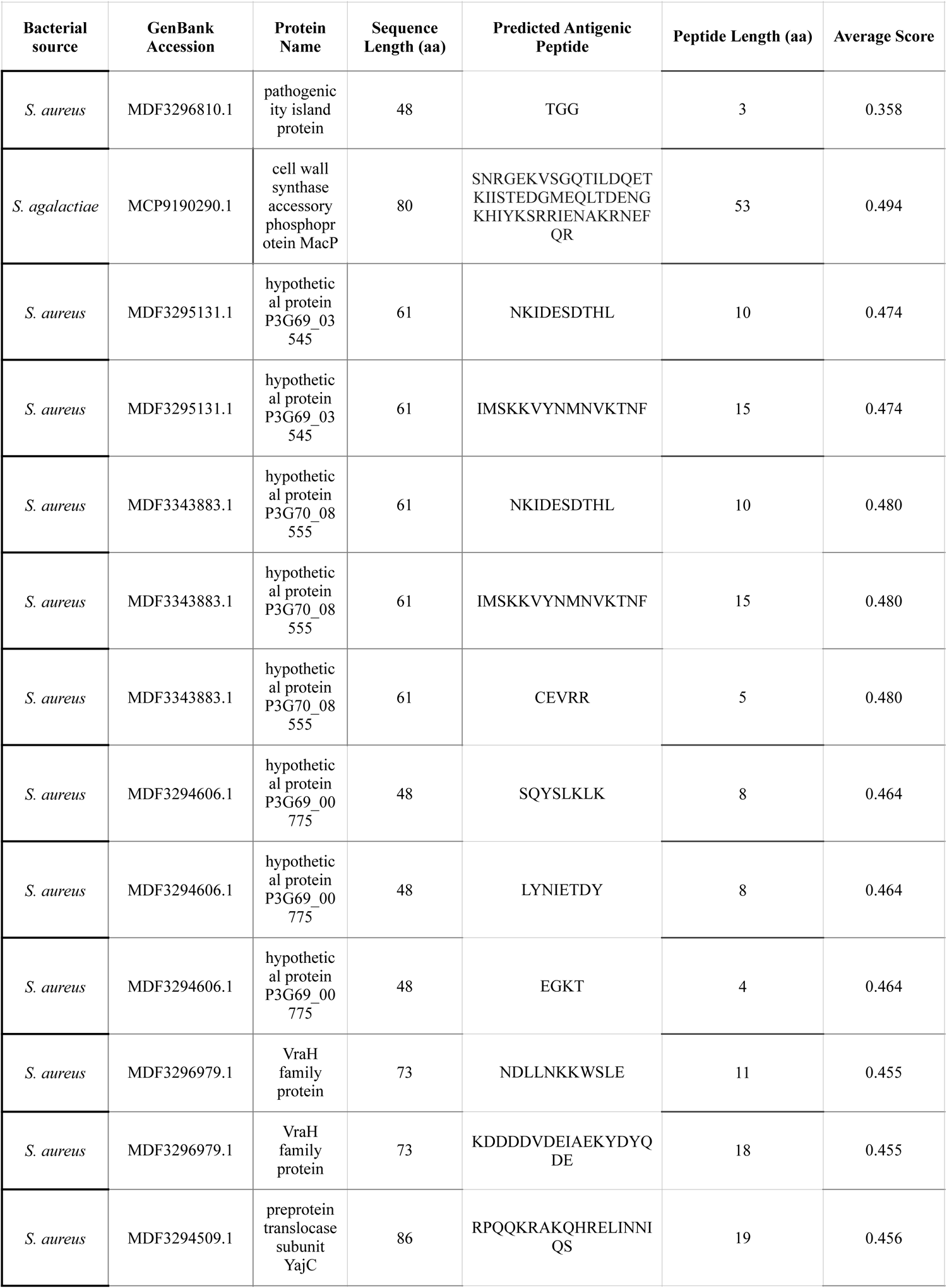
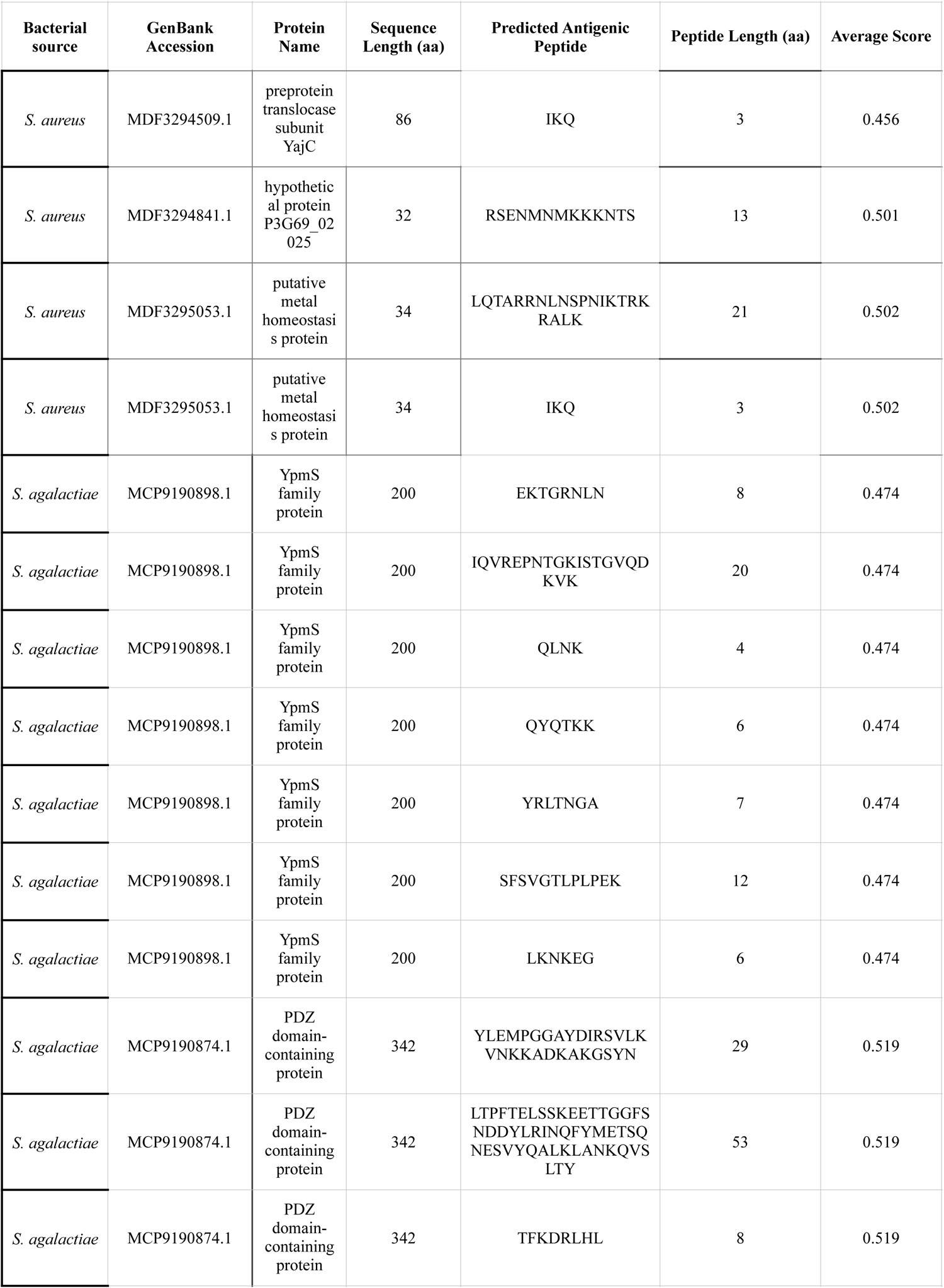
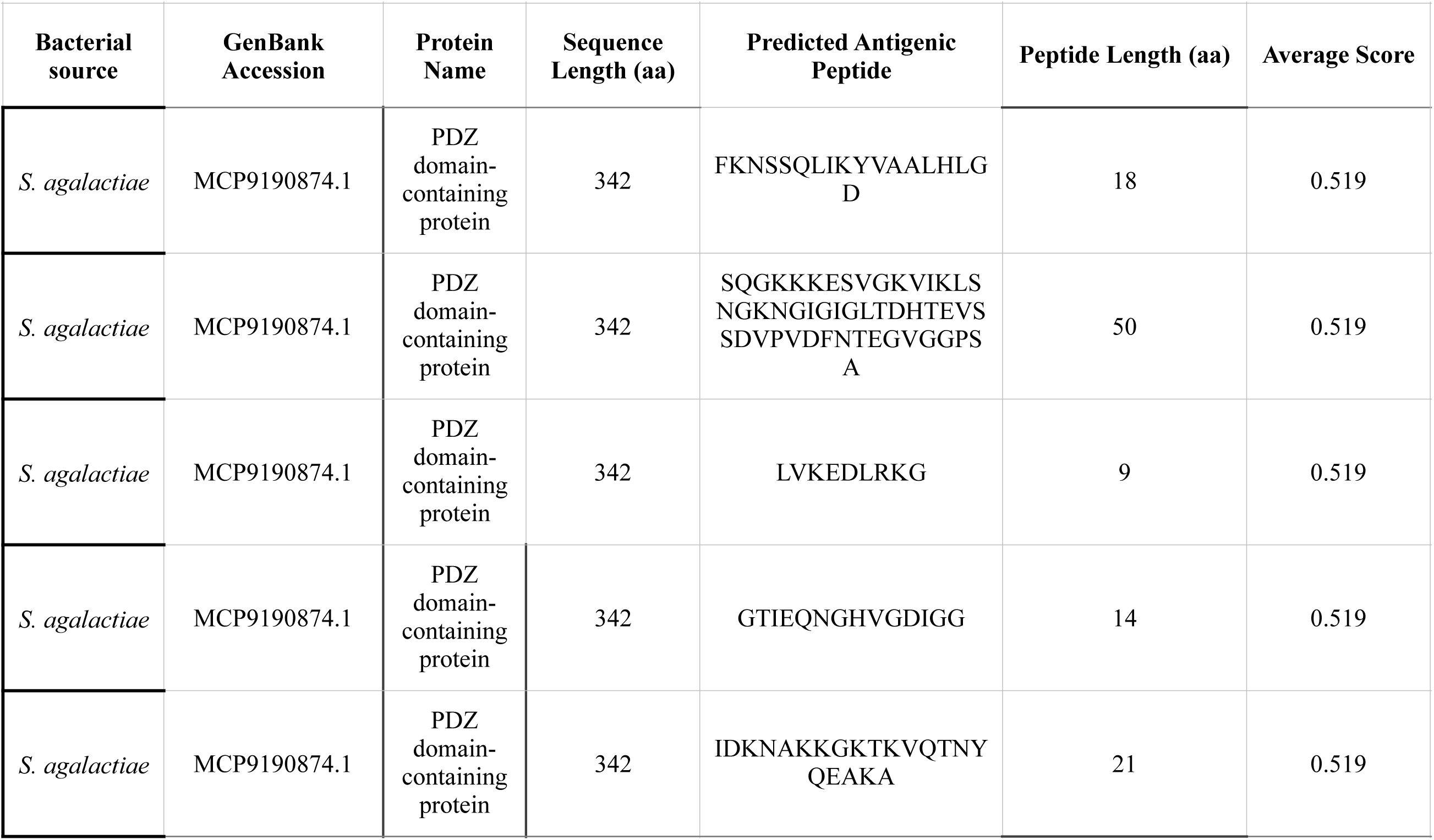
Predicted linear B cell epitope sequences in shortlisted *S. aureus* and *S. agalactiae* antigenic vaccine candidates. The length, composition and overall BepiPred antigenic score are shown.

#### 3.2.2 T cell (HTL and CTL) epitopes

MHC-1 molecules are expressed on the cell surface of all nucleated cells and serve to present peptide fragments derived from intracellular proteins to the immune system, alerting it to virally infected cells and subsequently triggering virus specific cytotoxic T lymphocytes (CTL) response (Hewitt, 2003). Collectively, 56 CTL were identified from several of the antigenic proteins under investigation using the NetCTL 1.2 server which predicts CTL epitopes in protein sequences (Table 3). Each predicted CTL epitope was assigned a particular score based on the prediction algorithm with only the top ten scoring motifs (indicating higher MHC I-binding affinity) considered for incorporation in the candidate vaccine following determination of their allergenicity and toxicity. Likewise, the NetMHCII 2.3 server was used for the prediction of 15-mer HTL epitopes from the selected antigenic proteins against a set of three human HLA alleles (HLA-DRB1*01:01, HLA-DRB1*01:03 and HLA-DRB1*03:01) (Supplementary Table 2). NetMHCII 2.3 output gives the affinity (nM) and strength of epitope binding (strong binding (SB) and weak binding (WB) to MHC-II molecules. HTL epitopes with strong binding (SB) to MHC-II molecules are shown in Table 4.

**Table 3:**
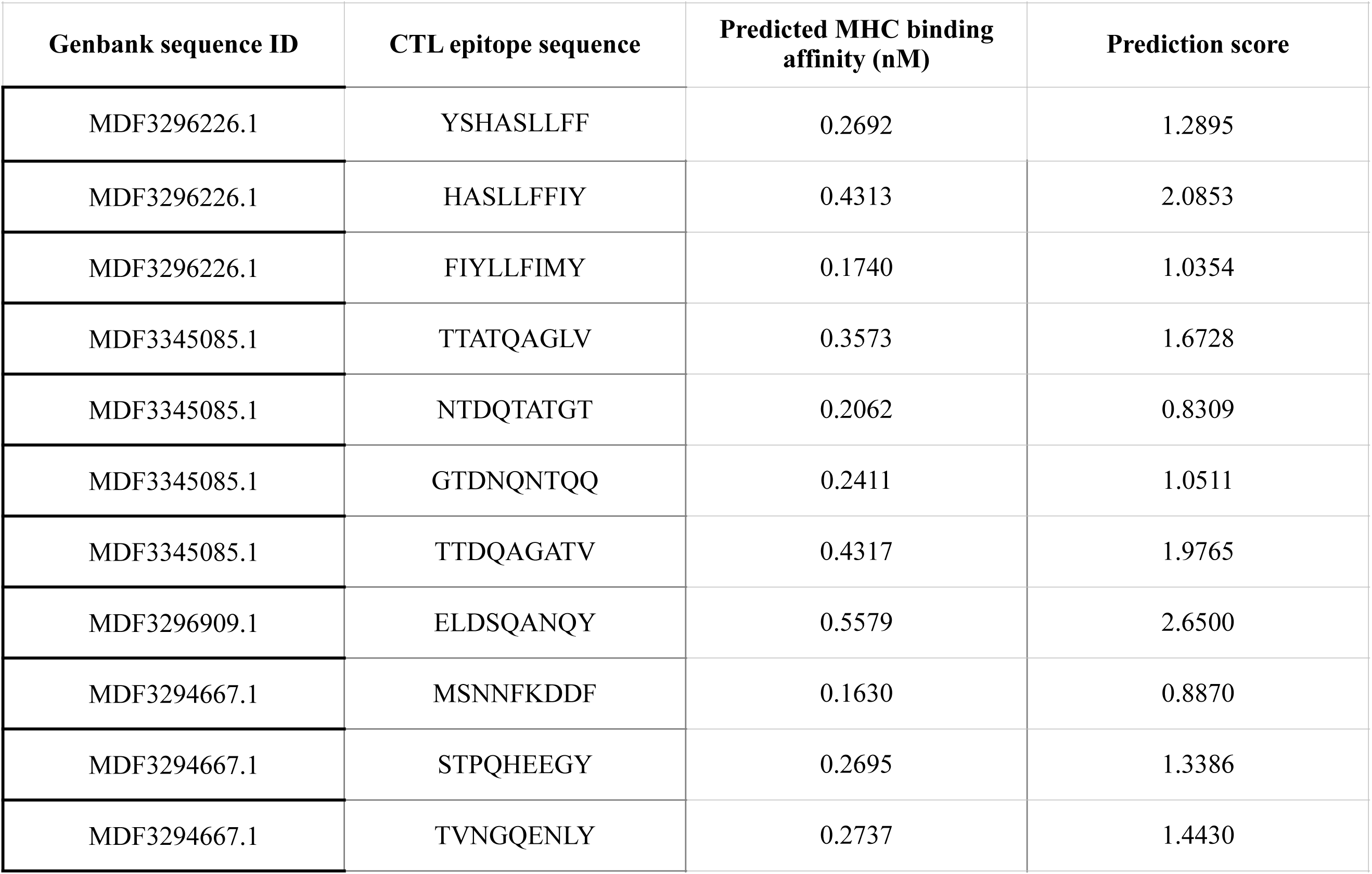
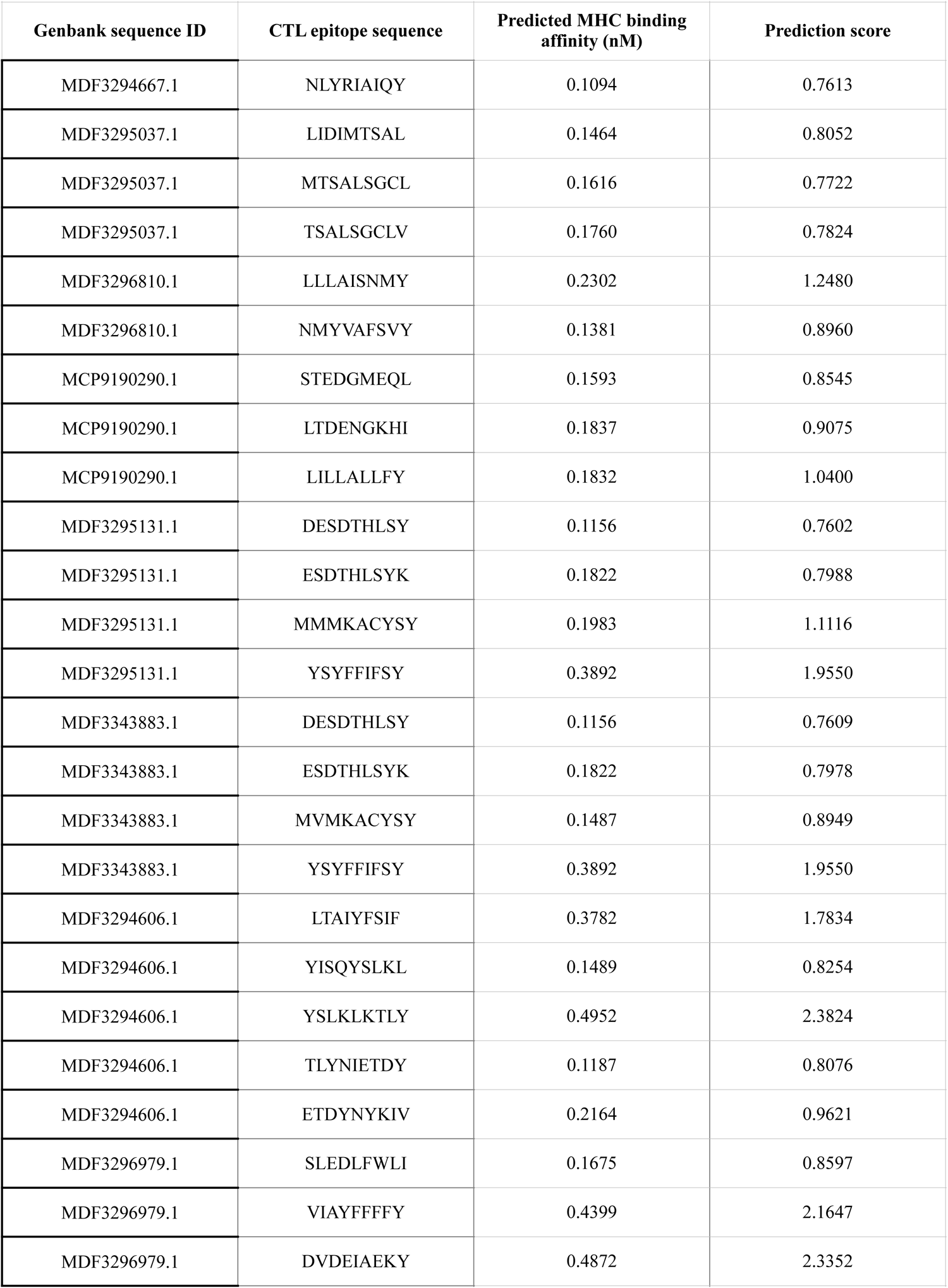
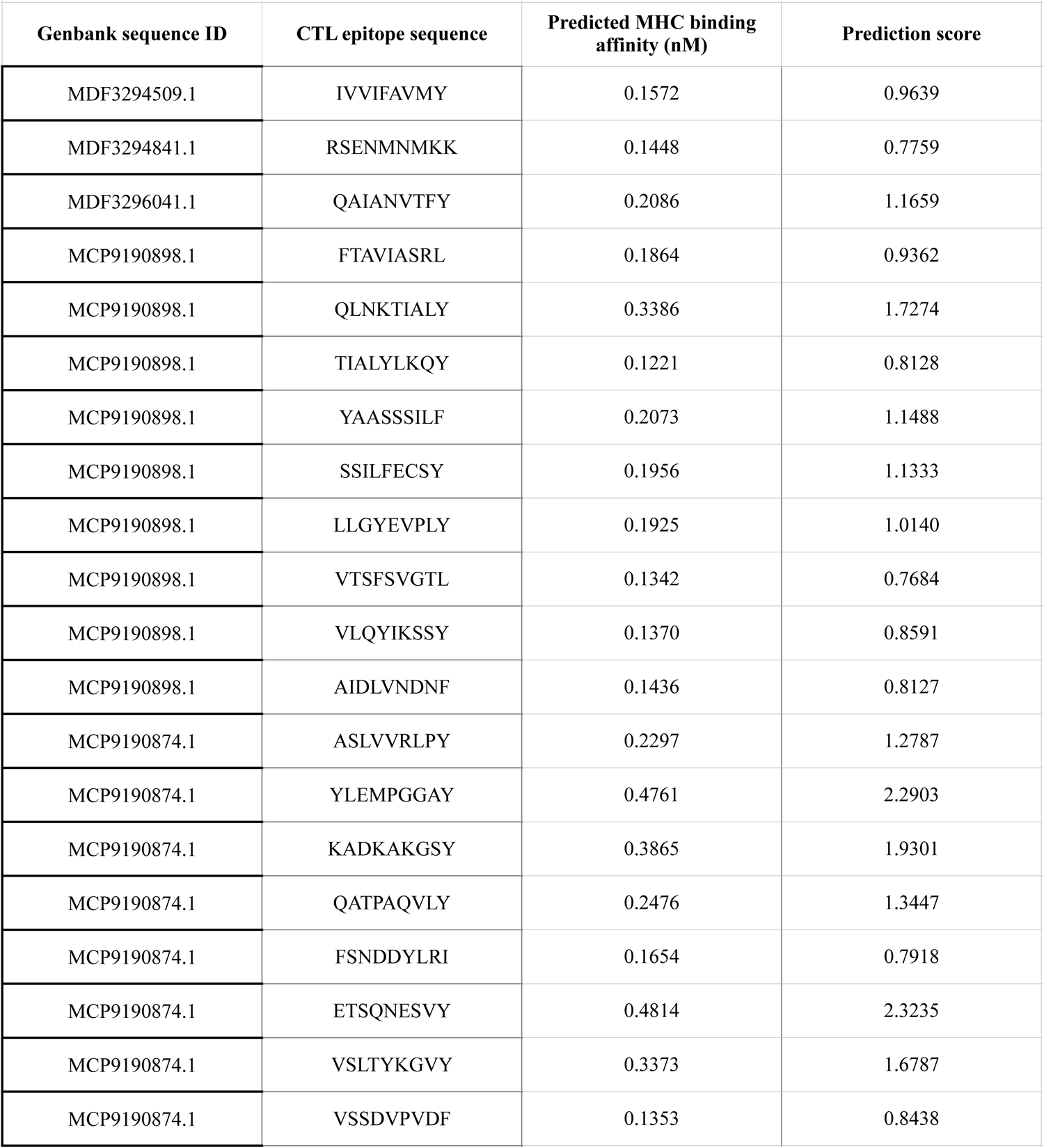
CTL epitopes predicted from the candidate vaccine sequences using the NetCTL 1.2 server.

**Table 4:**
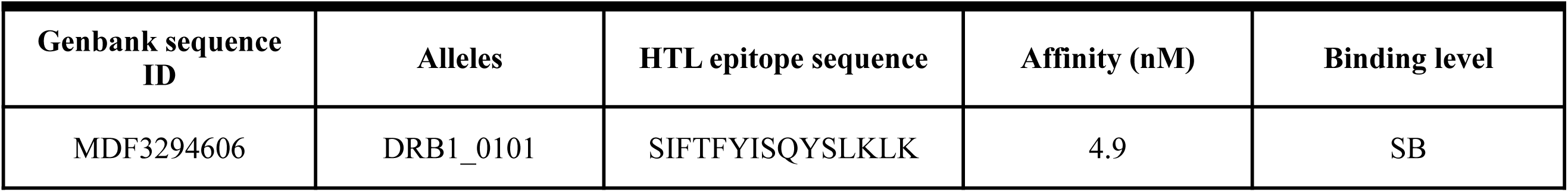
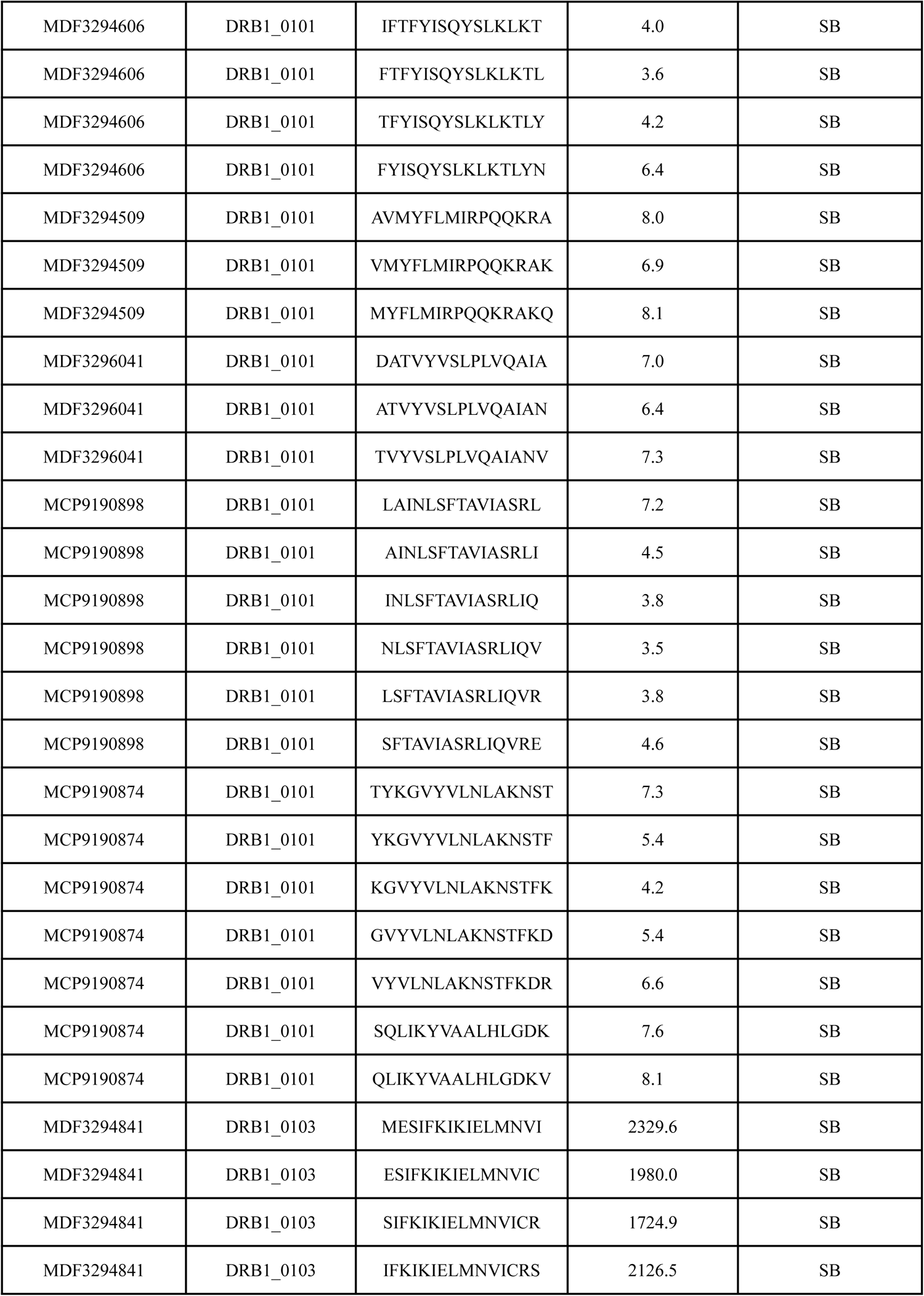
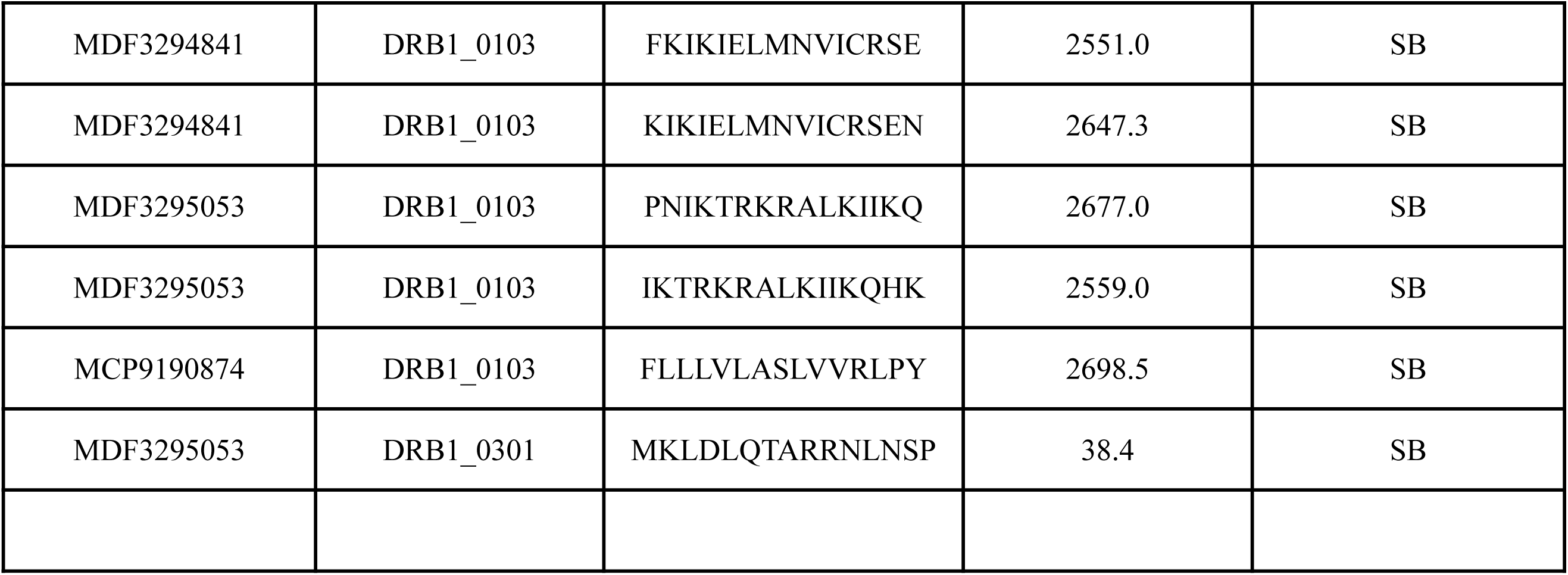
HTL epitopes predicted to possess strong binding (SB) to MHC-II molecules by the netMHCII 2.3 server. SB denotes strong binding.

The strongly binding HTL epitopes (Table 4) were evaluated for their ability to stimulate the production of interferon-γ using the IFNepitope server which predicts IFN-γ versus non-IFNγ epitopes using SVM and motif-based hybrid analysis. Seven peptides (Table 5) were revealed as potentially capable of triggering interferon-γ production. Thereafter, the seven peptides were assessed for their ability to stimulate IL-4 and IL-10 synthesis using the IL4pred and IL10pred servers, respectively. Both servers are underpinned by a SVM classifier that considers amino acid composition, dipeptide composition, amino acid propensity, and physicochemical properties as input parameters (Dhanda *et al.,* 2013). Indeed, the seven interferon-γ producing epitopes were predicted to trigger the production of IL-4 and IL-10 and thus appropriate for incorporation in the vaccine construct.

**Table 5:**
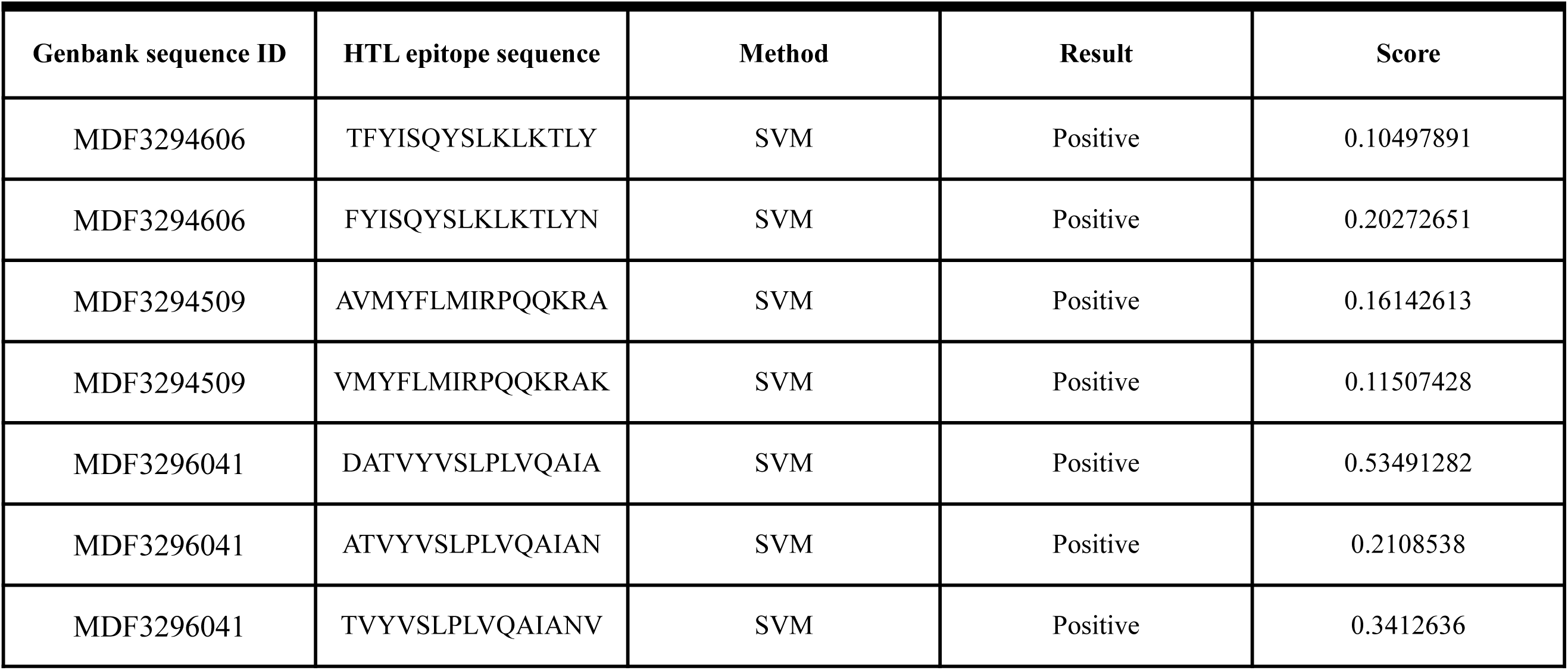
HTL epitopes with the potential of inducing interferon-γ production.

### 3.3 Construction of a candidate multi-epitope camel mastitis vaccine

The primary structure (amino acid sequence) of the multi-epitope vaccine was designed by combining the sequences of 9 CTL epitopes predicted to bind to MHC-I with high affinity and 1 interferon-γ inducing HTL epitope selected based in its MHC-II binding affinity and cytokine-inducing ability. The CTL and HTL epitopes were joined using using AAY and GPGPG linkers, respectively. To ensure that the vaccine candidates elicits a strong immune response, an adjuvant (219 amino acids long alpha chain of human IL-12; Uniprot ID: P29459) was added to the N-terminal of the construct using the EAAK linker (Afonso *et al.,* 1994; Bhattacharjee *et al.,* 2023). Moreover, to enable affinity purification following protein expression, a 6XHis tag was fused to the C-terminal of the vaccine construct. After the incorporation of the adjuvant, linkers and histidine tag, the final length of the candidate vaccine was 355 amino acids.

### 3.4 Evaluation of antigenicity, allergenicity, immunogenicity and toxicity

Determination of the antigenicity of the vaccine construct using VaxiJen returned a score of 0.5327 against a threshold of 0.4 stipulating that the candidate is potentially capable of eliciting host immune response. Evaluation of allergenicity using AlgPred 2.0 and AllerCatPro did not reveal any evidence of of sequence or structural similarity with known allergens indicating that the vaccine candidate is plausibly non-allergenic.

### 3.5 Physiochemical properties and solubility of the putative vaccine candidate

Using ProtParam, the average molecular weight of the vaccine construct was predicted to be 40074.76 with a theoretical isoelectric point (pI) of 6.04. Moreover, the candidate’s instability index (II) was 39.51, classifying it as a stable protein (II<40 indicates protein stability). The estimated *in vitro* half-life in mammalian reticulocytes was >30 h indicating *in vivo* stability of the putative vaccine candidate. The candidate had an aliphatic index and grand average of hydropathicity (GRAVY) of 86.74 and −0.097 respectively, indicating that thermostability and aqueous solubility.

### 3.6 Secondary and tertiary structure features of the vaccine candidate

The 3-D model of the putative vaccine construct inferred using I-TASSER revealed it to contain 8 helices and 2 beta strands (Figure 2). Analysis os the model’s quality using ProSA revealed it to be robust with a Z-Score of −4.16.

**Figure 1:**
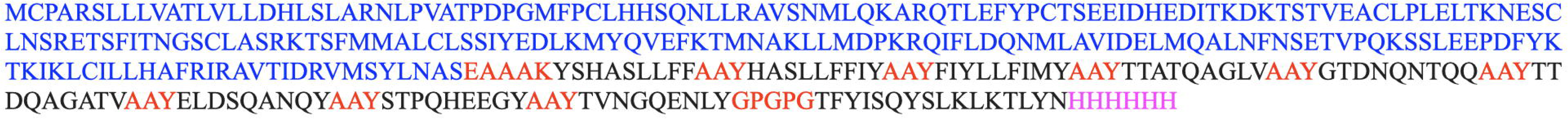
Amino acid sequence of the candidate multi-epitope vaccine. The vaccine adjuvant sequence is depicted in blue, CTL and HTL epitopes are rendered in black, linkers (EAAAK, AAY and GPGPG) are shown in red while the His-tag is illustrated in magenta.

**Figure 2:**
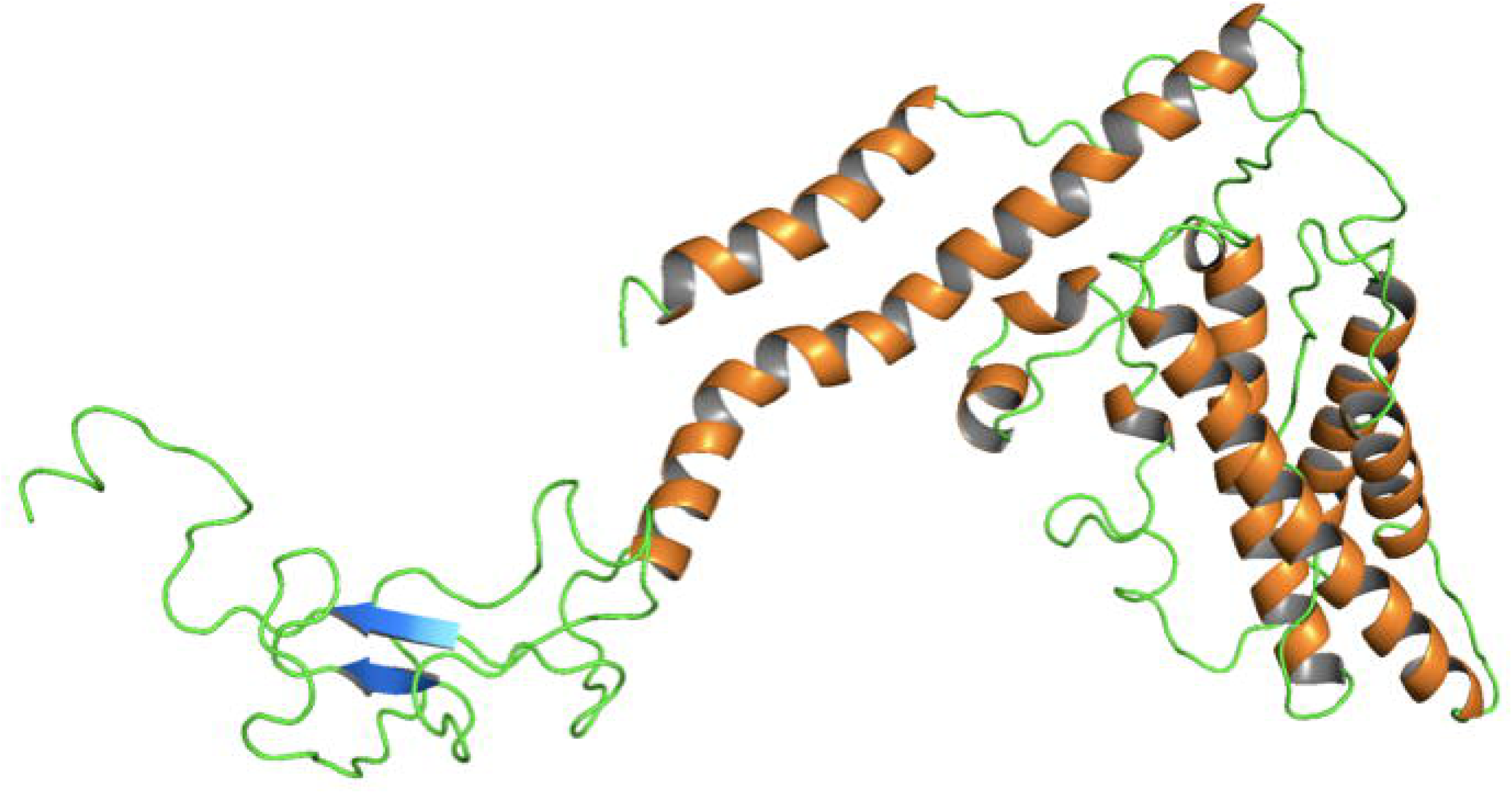
Cartoon representation of the predicted three-dimensional (3-D) structure of the vaccine construct. Helices, beta sheets and loops are coloured orange, magenta and green respectively.

### 3.7 Molecular docking of the vaccine construct onto TLR2

The CASTp server identified various binding sites in the vaccine construct with the largest having a binding area and volume of 326.094 Å2 and 244.195 Å3 respectively (Supplementary Table 3). However, ClusPro server predicted the interaction with TLR-2 (−1265.1 kJ/ml) to plausibly occur via a much smaller pocket in the C-terminal region of the construct (Figure 3). Activation of the membrane-bound receptor TLR-2, the most indiscriminate of the TLRs with respect to bacteria, virus, parasites and fungi derived pathogen-associated molecular patterns (PAMPs), triggers the production of nuclear factor-kappa B and cytokine and subsequent stimulation of innate immunity (Akira *et al.,* 2006).

**Figure 3:**
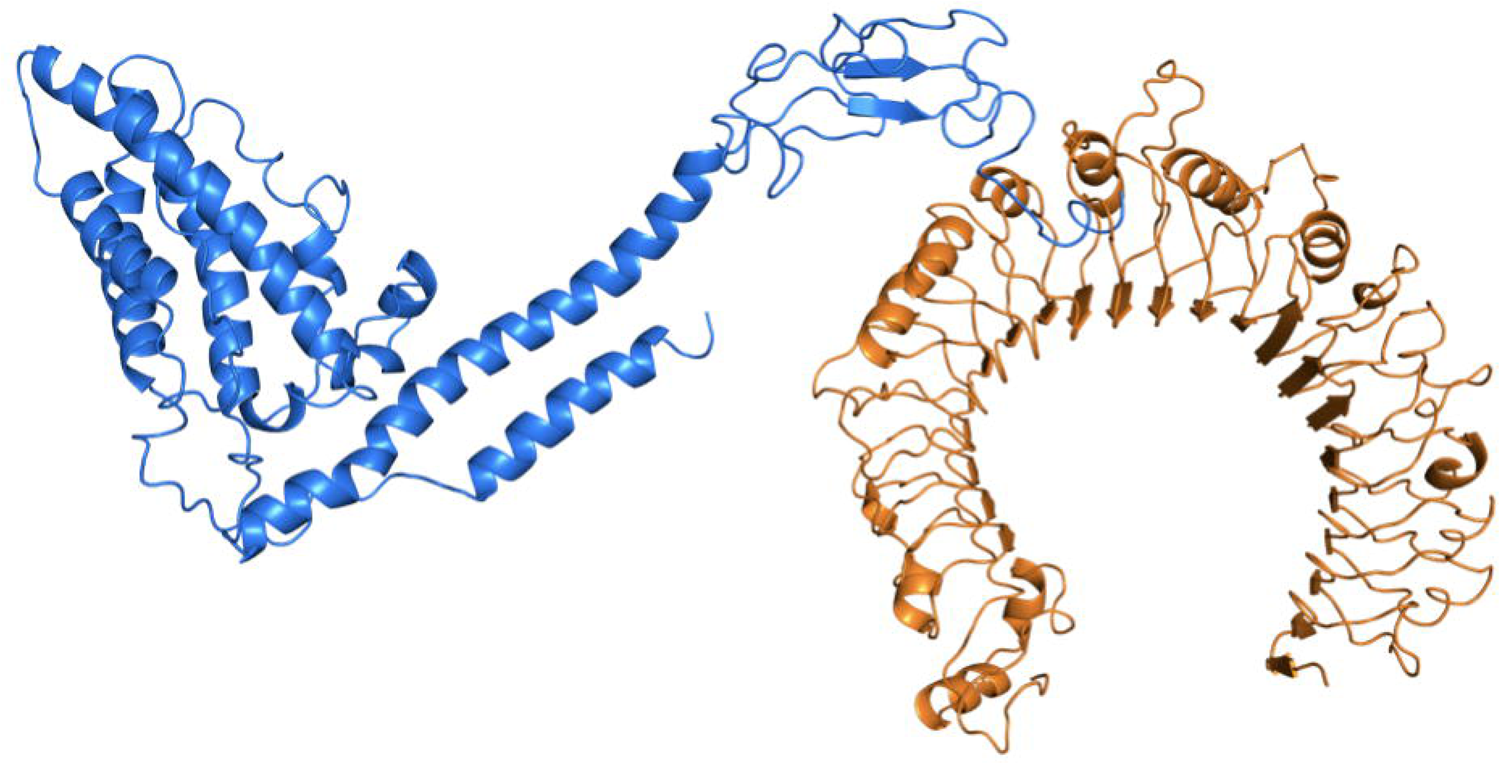
Depiction of the vaccine construct (marine) and TLR-2 (orange) complex.

### 3.8 Molecular dynamics (MD) simulation of the vaccine candidate-TLR2 complex

The iMODS server was used to explore the collective motions of the vaccine construct-TLR-2 complex (Figure 4). The overall Eigenvalue of the complex was 5.155235e-07. The deformability plot (Figure 4A) indicates that the complex is adequately flexible.

**Figure 4:**
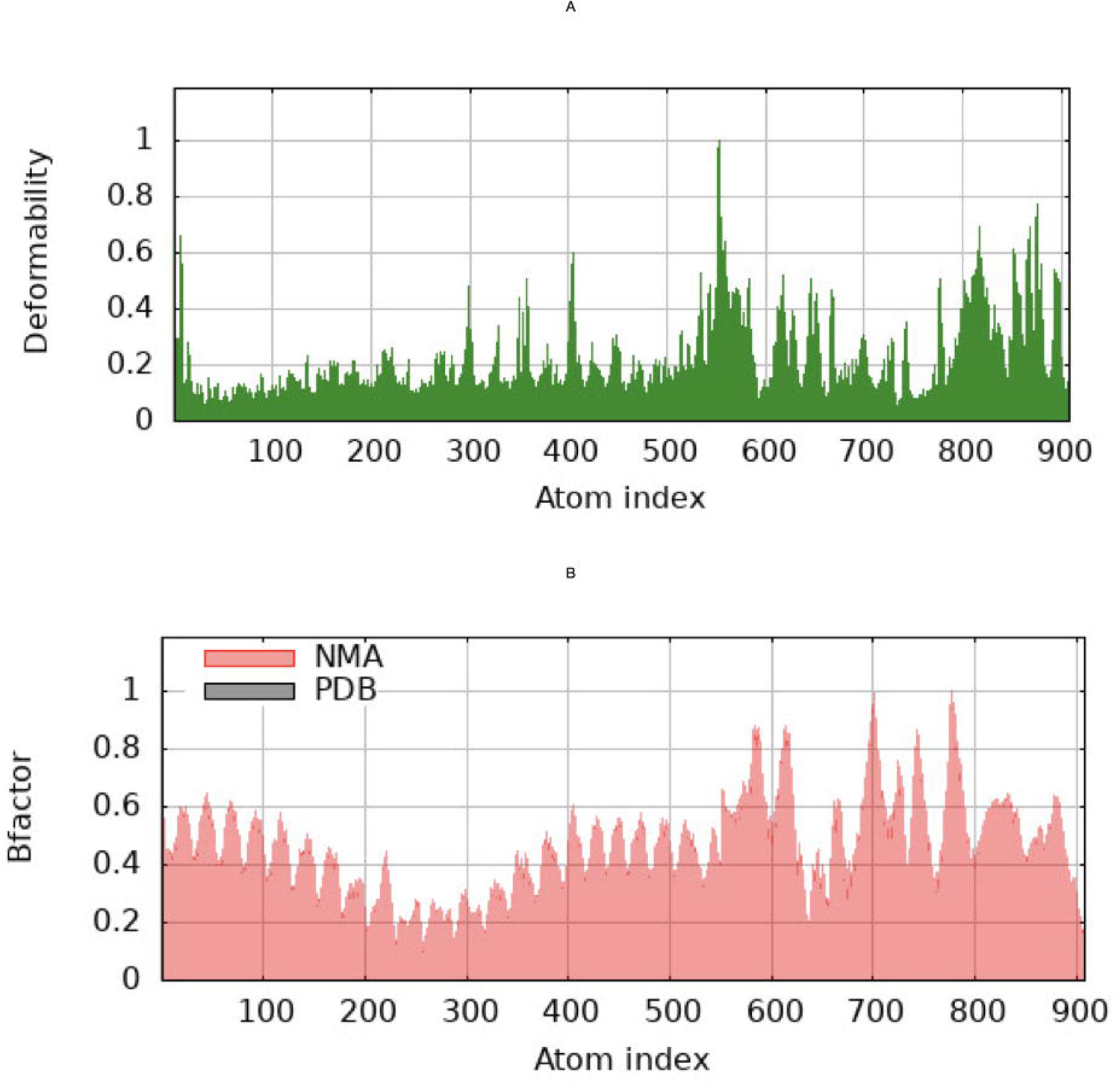
An illustration of deformability **(A)** and B-factor **(B)** of the vaccine construct and TLR-2 complex ascertained using the iMODS server. The main-chain deformability is a measure of the ability of a given molecule to deform at each of its residues. High deformability regions indicate chain ‘hinges’.

### 3.9 Simulation of the ability of the vaccine construct to elicit host immune response

The vaccine construct was subjected to *in silico* immune simulations to ascertain its potential to stimulate the adaptive immune system. The results obtained indicate that it is capable of inducing production of high levels of IgG and IgM with a corresponding reduction in antigen levels (Figure 5). Additionally, it was shown to be capable of triggering synthesis of T-helper lymphocytes, T-cytotoxic lymphocytes and, cytokines and interleukins In sum, the designed vaccine construct would potentially elicit a strong immune response.

**Figure 5:**
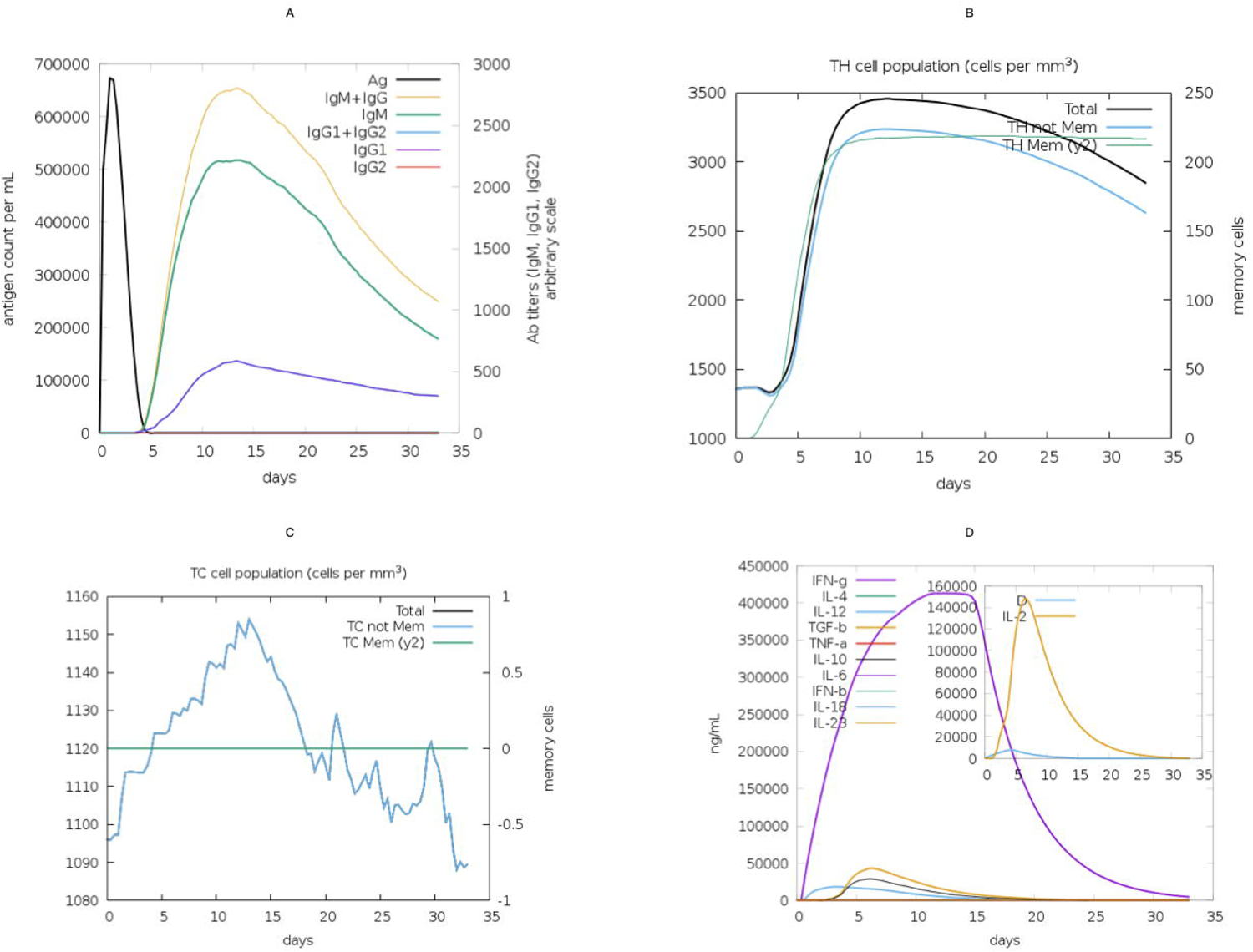
Depiction of the simulated ability of the vaccine construct to trigger antibody production **(A)**, T-helper lymphocytes **(B)**, T-cytotoxic lymphocytes **(C)**, cytokines and interleukins **(D)**.

### 4.0 Discussion

Camel mastitis (inflammation of the mammary gland) causes immense socio-economic losses for pastoral and nomadic communities in the arid and semi-arid (ASALs) regions of Africa where the one-humped camels (*Camelus dromedarius*) are an invaluable source of livelihood and nutrition due to their resilience to severe droughts that cause high cattle, sheep and goat mortality (Mwangi *et al.,* 2022). Specifically in the ASALs of Kenya, camel productivity is severely threatened by severe droughts linked to climate change and mastitis which endanger the livelihoods and nutrition of pastoralists in these regions.

The importance of camels in buttressing economic and food security for pastoralists in these regions is bound to become increasingly entrenched by the confluence of the adverse impacts of climate change and extensive dissemination of antimicrobial resistance (AMR) that cause frequent severe droughts and render current antimicrobials largely ineffective respectively.

Notably, unlike in other dairy animals where treatment of mastitis with intra mammary antibiotics is effective, this approach is very challenging in camels due to the unique anatomy of camelidae udder (Aqib *et al.,* 2018). Therefore, there is a pressing need for the development of a novel, efficacious camel mastitis vaccine. Indeed, the Kenya AMR Policy and AMR Action Plan recommend the use of vaccines as a sustainable disease control option. In the One-Health framework, reverse vaccinology is a particularly attractive strategy for accelerating the development of human and livestock vaccines in resource limited regions given its ability to significantly shorten development timelines and thus lower cost (Dzayee *et al.,* 2022). Specifically, using this methodology, potential immunogenic, non-allergenic, non-toxic candidates are rapidly predicted and prioritised for experimental evaluation.

In this study, we have designed and computationally evaluated a novel, multi-epitope vaccine candidate for camel mastitis comprised of 9 CTL MHC-1 binding epitopes and 1 interferon-γ inducing HTL epitope obtained from selected antigenic protein sequences of local isolates of *S. agalactiae* and S. *aureus* (Murungi *et al.,* 2022; Maichomo *et al.,* 2023), the predominant camel mastitis causing bacteria (Younan *et al.,* 2001; Seligsohn *et al.,* 2021a; Seligsohn *et al.,* 2021b). *In silico* simulations revealed the vaccine construct as capable of strongly stimulating the host innate and adaptive immune responses as evidenced by the elevated synthesis of IgM, IgG, cytokines, macrophages, dendritic cells and natural killer cells. The antigenic proteins from which the epitopes were derived are largely cytoplasmic membrane proteins. Surface immunogenic proteins have been demonstrated to be potent inducers of immune responses against microbial infections (Diaz-Dinamarca *et al.,* 2020). For example, iron-regulated surface protein A (IsdA) and surface immunogenic protein (Sip) have been proposed as vaccine candidates for bovine mastitis vaccine (Chatterjee *et al.,* 2021; Pathak *et al.,* 2022). The antigenic proteins from which the epitopes used in the design of the candidate vaccine include *S. aureus* elastin binding protein (EbpS), type I toxin-antitoxin system Fst family toxin, pathogenicity island protein, cell wall synthase accessory phosphoprotein MacP, VraH family protein, preprotein translocase subunit YajC, YpmS family protein and PDZ domain-containing protein. EbpS is an integral membrane protein that plays a key role in the virulence of *S. aureus* together with other factors including cytolytic toxins, superantigens and extracellular proteins (Downer *et al.,* 2002). Specifically, EbpS is an adhesin that binds to the host extracellular matrix component elastin (Campoccia *et al.,* 2009). Another candidate protein, type I toxin-antitoxin system Fst family toxin, detoxifies toxins produced by bacteria and by so doing ensures that bacterial replication, translation and cell division proceed (Masachis and Darfeuille, 2018). On the other hand, the membrane-anchored cell wall synthase accessory phosphoprotein MacP, a cofactor of PBP2a (a penicillin binding protein (PBP) synthase) has been been shown to activate the cell wall synthase PBP2a of *Streptococcus pneumoniae* in a phosphorylation-dependent manner (Fenton *et al.,* 2018) thus contributing to cell growth and division. The small staphylococcal transmembrane protein VraH forms, together with VraDE, a three-component system that plays a role in antibiotic resistance and pathogenicity (Popella *et al.,* 2016).

Given that cell-mediated immune response is the principal mode through which the host counters mastitis, the designed vaccine candidate comprised of 9 CTL MHC-1 binding epitopes and 1 interferon-γ inducing HTL epitope. Linking of the CTL epitopes was via AAY while the HTL epitope was linked using a GPGPG linker. Moreover an adjuvant (alpha chain of human IL-12) was incorporated at the N-terminal end of the construct. The construct is predicted to be non-allergenic, non-toxic, highly stable and soluble. PRoSA evaluation revealed the the 3-D model structure of the designed vaccine construct to be of robust quality. Moreover, the model was shown to be adequately flexible therefore capable of interaction with the immune system components. *In silico* immune simulations indicated the potential vaccine candidate as not only capable of inducing production of high levels of IgG and IgM with a corresponding reduction in antigen levels but also triggers synthesis of T-helper lymphocytes, T-cytotoxic lymphocytes and, cytokines and interleukins.

## 5.0 Conclusion

Collectively, we have designed and computationally evaluated a candidate vaccine for camel mastitis that exhibits desirable properties (immunogenic, non-allergenic and non-toxic) and warrants further experimental investigations. Our findings represent a significant step in the camel mastitis vaccine development endeavours.

## Data availability statement

The original contributions presented in this study are included in the article/Supplementary Material. Further inquiries can be directed to the corresponding author.

## Author Contributions

Conceptualisation, E.M., N.G., H.W., M.M.; methodology, E.M., I.O., E.M., N.L., R.O.; software, E.M., I.O., E.M., N.L., R.O.; validation, E.M., N.L., R.O.; formal analysis, E.M., E.M., N.L., R.O.; investigation, E.M., E.M., N.L., R.O.; resources, H.W., M.M.; writing—original draft preparation, E.M., E.M., N.L., H.W., M.M.; writing—review and editing, E.M.; visualisation, E.M., E.M., N.L., R.O.; supervision, E.M., H.W., M.M. All authors have read and agreed to the published version of the manuscript.

## Acknowledgements

We thank AGRI-FI and KALRO for financial and technical assistance respectively.

## Publisher’s note

All claims expressed in this article are solely those of the authors and do not necessarily represent those of their affiliated organizations, or those of the publisher, the editors and the reviewers. Any product that may be evaluated in this article, or claim that may be made by its manufacturer, is not guaranteed or endorsed by the publisher.

## Supplementary material

The Supplementary Material for this article is available online.

